# Spatially clustered neurons encode vocalization categories in the bat midbrain

**DOI:** 10.1101/2023.06.14.545029

**Authors:** Jennifer Lawlor, Melville J. Wohlgemuth, Cynthia F. Moss, Kishore V. Kuchibhotla

**Affiliations:** Department of Psychological and Brain Sciences, Johns Hopkins University, Baltimore, MD, 21218, USA; Johns Hopkins Kavli Neuroscience Discovery Institute, Baltimore, MD, 21218,MD; Department of Neuroscience, University of Arizona, Tucson, AZ, 85721, USA; The Solomon Snyder Department of Neuroscience, Johns Hopkins University School of Medicine, Baltimore, MD, 21205, USA; Department of Mechanical Engineering, Johns Hopkins University, Baltimore, MD, 21218, USA; Department of Biomedical Engineering, Johns Hopkins University, Baltimore, MD, 21218, USA

## Abstract

Rapid categorization of vocalizations enables adaptive behavior across species. While categorical perception is thought to arise in the neocortex, humans and other animals could benefit from functional organization of ethologically-relevant sounds at earlier stages in the auditory hierarchy. Here, we developed two-photon calcium imaging in the awake echolocating bat (*Eptesicus fuscus)* to study encoding of sound meaning in the Inferior Colliculus, which is as few as two synapses from the inner ear. Echolocating bats produce and interpret frequency sweep-based vocalizations for social communication and navigation. Auditory playback experiments demonstrated that individual neurons responded selectively to social or navigation calls, enabling robust population-level decoding across categories. Strikingly, category-selective neurons formed spatial clusters, independent of tonotopy within the IC. These findings support a revised view of categorical processing in which specified channels for ethologically-relevant sounds are spatially segregated early in the auditory hierarchy, enabling rapid subcortical organization of call meaning.

## Introduction

Vocal communication exemplifies the ecological importance of categorical perception. Humans readily recognize the identity of a single person despite similarity in spectrotemporal features among many individuals. Moreover, categorical discrimination of vocal communication calls appears to be phylogenetically conserved across species, including those that communicate on land (non-human primates^1^, naked mole rats^2^, mice^3^, crickets^4^, frogs^5^), sea (dolphins^6^, pinnipeds^7^), and air (songbirds^8^, echolocating bats^9^). Current models of categorical perception propose that sensory cortical networks are the engine underlying category formation^10^ – binding relevant features across a basis set of lower-level features that are initially decomposed from natural scenes by sensory organs and processed hierarchically in a feedforward manner ^11–13^. While the perception of categories may arise in the cortex, categorical representations at earlier stages in sensory processing would provide significant computational advantages, conferring increased speed and fidelity of categorization.

To what extent do categorical representations emerge earlier in the sensory hierarchy? Categorical representations may exist in the form of single neuron selectivity, population selectivity, or even in spatially segregated ensembles^14^. We reasoned that applying an optical approach, which provides high spatiotemporal resolution, in a mammalian species that exhibits a rich repertoire of vocalization categories would allow us to test these possibilities. The big brown bat, *Eptesicus fuscus*, is an insectivorous bat species that stands out in the animal kingdom for its acoustic repertoire for echolocation-based navigation^15^ and for social interactions with conspecifics^16,17^. *E. fuscus* moves in three dimensions through its environment and must rapidly distinguish between vocalizations produced for navigation or social interaction to successfully hunt and communicate.

## Results

### Two-photon calcium imaging in the awake, echolocating bat

One candidate subcortical region is the inferior colliculus (IC) because of its key position as an obligatory auditory processing station in the central nervous system. It has multiple subdivisions that receive inputs from the auditory brainstem ^18^. One subdivision, the dorsal cortex of the IC (DCIC), is of particular interest for categorical representations as it receives both feedforward input and significant top-down input from the auditory cortex^19,20^. Here, we developed an approach for two-photon imaging in the inferior colliculus (IC) of the awake, big brown bat, *E. fuscus* (**Figure 1**) to study spatially resolved single neuron and population coding of auditory stimuli. We first sought to characterize the sound-evoked properties of DCIC neurons and validate the use of two-photon calcium imaging in this non-traditional animal model. We injected AAV5-CAMKII-GCaMP6f bilaterally into the DCIC of *E. fuscus* and used a thinned-skull procedure to gain chronic optical access (**Figure 1A-B**). We used two-photon imaging to monitor large populations of excitatory neurons (**Figure S1A-B**) with single-cell resolution (**Figure 1C**). We imaged from multiple sites in the left and right IC (LIC and RIC, **Figure 1B**), and at multiple depths, from the brain’s surface and up to 200 *μ*m. Using this technique, we recorded neural activity from 7,868 spatially-resolved neurons across 20 sites and 3 bats. Awake bats were presented with an array of auditory stimuli including pure tones (4-100 kHz), complex stimuli (FM sweeps and white noise), and a series of bat and mouse vocalizations. Individual neurons exhibited robust sound-evoked responses (**Figure 1C-D**) with most of the recorded population responding to at least one of the stimuli presented (**Figure 1E-F**, n=3 bats, 79.6±3.7%, p < 0.05, paired t-test).

**Figure 1.**
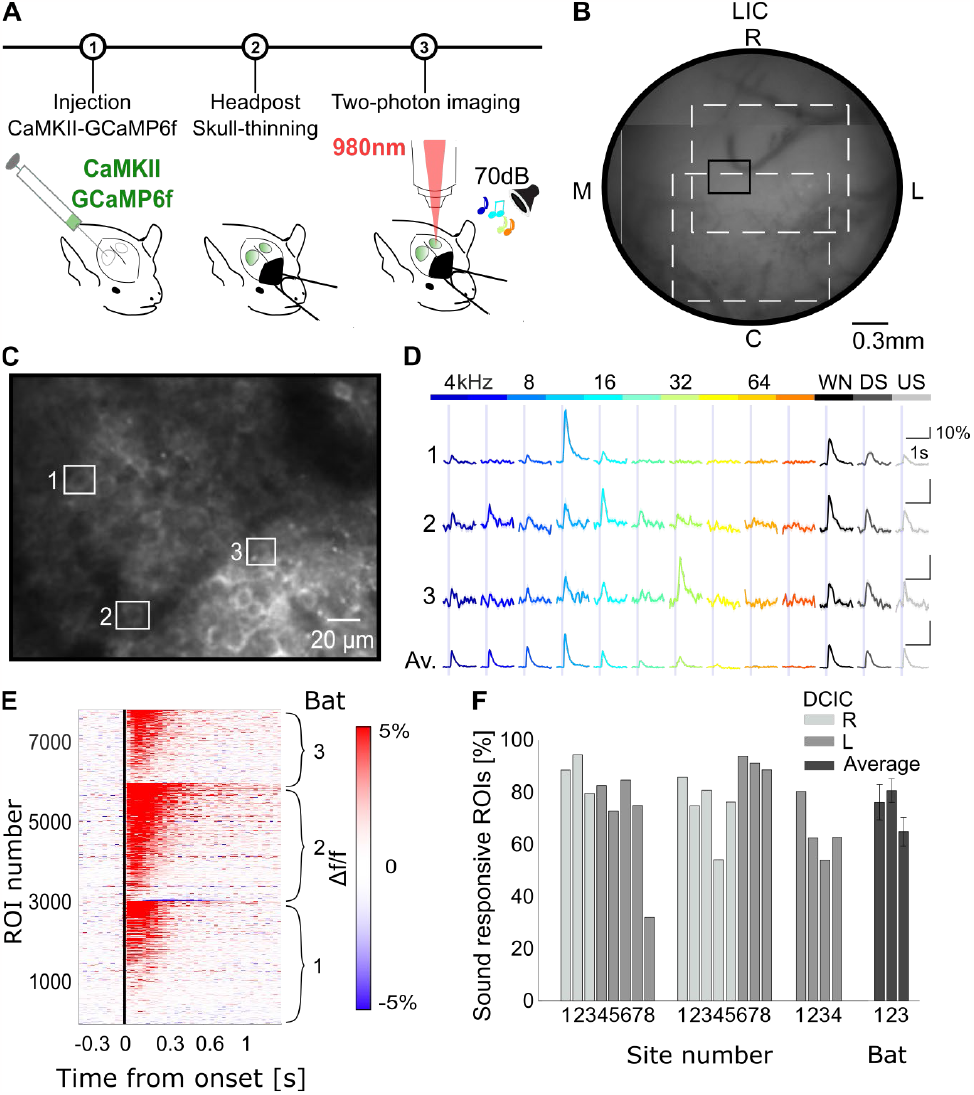
Robust sound-evoked responses in the DCIC of awake bats observed using two-photon calcium imaging. **A**. In vivo two-photon calcium imaging methods and timeline graphical representation. **B**. Composite vasculature map of an example LIC with overlaid imaging sites at 2X magnification (white box). The small black box highlights the position of the cells displayed in C. **C** Example site mean image cropped. White boxes surround examples cells presented in D. **D**. Average activity (Δf/f) of 3 example ROIs to pure tones and complex sounds presentations (color indicates stimulus identity displayed above, shading: SEM, WN: white noise, DS: frequency modulated down sweep, US: frequency modulated up sweep). The fourth calcium trace is the average trace for the imaged population of the example site (Av., shading: SEM). **E**. Sound-evoked heatmap for all identified ROIs across all sites and bats (n_cells_ = 7868, averaged over all stimuli types presented in this manuscript). **F**. Percentage of significantly sound-evoked ROIs per site (light grey: right DCIC, medium grey: left DCIC) and averaged across site (darker grey) for each bat.

### Functional microarchitecture of the DCIC

We next sought to understand the functional microarchitecture of the DCIC in echolocating bats. A fundamental feature of the auditory system is the precise topographic organization of sound frequency, commonly referred to as ‘tonotopy’. Tonotopy emerges from the biomechanical properties of the basilar membrane in the inner ear and propagates throughout the central auditory system^21^. The central nucleus of the IC (CNIC) in bats and other mammalian species exhibits a dorso-ventral tonotopic axis^22^ but less is known about the topographical organization of sound frequency in the DCIC, in part because prior studies in the bat have not consistently identified the precise recording site ^23,24^. We presented pure tones ranging from 4 to 90 kHz, matching *E. fuscus’* hearing range^25^, at 70 dB with a half-octave spacing and then calculated single-neuron tuning curves to identify a neuron’s ‘Best Frequency’ (BF), here defined as the sound frequency that elicits the highest amplitude stimulus-evoked response. Most neurons were significantly tuned to a limited band of sound frequencies (66.4±11%, two-way ANOVA for tone and duration, p<0.05, **Figure 2A-C**) in the low-to-mid frequency range (**Figure 2C**, 4 to 32 kHz: 93.3%±1.7% of tuned neurons). These findings are consistent with those reported in the mouse using two-photon calcium imaging^19,26^. In addition, these data are consistent with anatomical evidence showing that antero-ventral cochlear nucleus neurons that encode lower-frequency sounds project more superficially than dorsal axonal projections^27^, suggesting a conserved functional microarchitecture across species.

**Figure 2.**
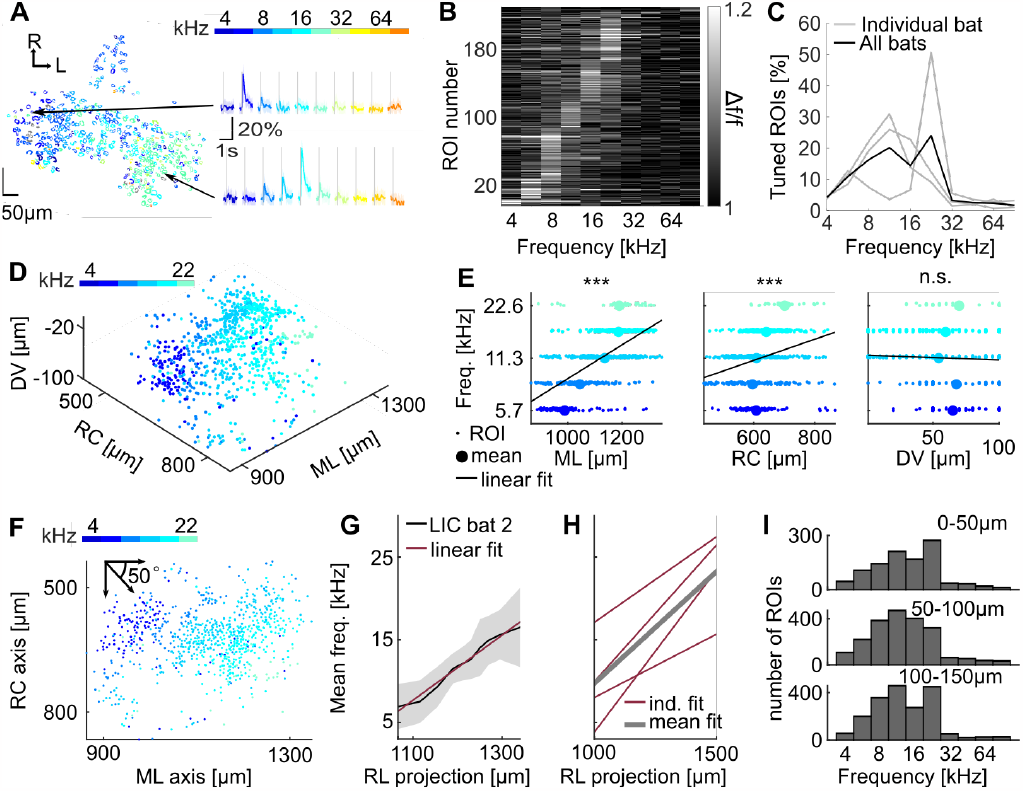
Superficial tonotopy in the DCIC. **A**. Left: example site from the RIC of bat 2, ROIs color-coded by their BF. Right: tone-evoked PSTHs for example cells. The first cell showed clear tuning to 5.7 kHz and the second to 16 kHz. **B**. Tuning curves for all tone-responsive ROIs from the example site presented in A. **C**. Distribution of tuning per bat (grey, n = 3) and averaged (black) as a percentage of the tuned population. Most of the ROIs’ tuning is in the low to mid-frequency range (93.3±1.7% of tuned neurons’ BF between 5.7 to 22.6kHz) **D**. 3D reconstruction of bat 2 LIC (n_site_ = 3, n_cells_= 835), tuned ROIs are color-coded according to their BF, showing low frequency rostrally and medially and higher frequency caudally and laterally **E**. ROIs’ BF (individual ROI: small dot, average: larger dot) as a function of distance along three along three anatomically defined axis for example LIC in D for each frequency (left: ML, middle: RC and right: DV). Linear fit (black line) is significant for ML and RC but not for DV (p=5.02e-95, p=4.79e-15 and p =0.13, respectively) **F**. 2D reconstruction of bat 2 LIC, tuned ROIs are color-coded according to their BF showing frequency bands oriented rostromedially. **G**. Mean BF (in black) and its linear fit (in red) along the 50° RL axis for LIC in F. Mean tuning increases along this axis, confirming the existence of a tonotopic gradient. **H**. Linear fit (in red) of tuning along the 50° RL axis for each bat hemisphere (in red) and the average (in grey). Tuning increases as a function of RL distance by 2.1oct./mm on average. **I**. Frequency tuning distribution across bats along the DV axis (50 *μ*m bins from 0 to 150 *μ*m). Tuning does not significantly vary with depth (Friedman test, p_dv_=0.4066).

We then used the spatial resolution of two photon imaging to reconstruct the precise location of each neuron in three dimensions with its corresponding BF (**Figure 2D**). We aligned across imaging sites and bats in the rostrocaudal and the mediolateral extents, based on a common anatomical landmark, the intersection of the IC, superior colliculus and midline. Z-depth was measured using the optical sectioning capabilities of two-photon imaging coupled with an estimate of the skull-to-brain transition, based on a change in background fluorescence (**Figure S2A-C**). This allowed us to test whether a spatial, tonotopic gradient exists in the mediolateral, rostrocaudal, or dorsoventral axes. We determined that a positive linear relationship exists for both the mediolateral and the rostrocaudal axes but not for the dorsoventral axis (**Figure 2E**, p_ML_ = 5.02e^-95^, p_RC_= 4.79e^-15^, and p_DV_ = 0.13, additional example sites in **Figure S3A**). To capture the direction of this tonotopic axis, we estimated the best fit angle per reconstructed hemisphere across bats (**Table S1**), yielding an average angle of 47.5°, a value remarkably close to the 50° reported in the mouse DCIC ^26^. The tonotopic gradient was computed as the linear fit of the mean frequency along this rostrolateral axis (**Figure 2F-G**) per hemisphere and per bat. On average, the BF increased by 2.1 oct./mm along this axis (**Figure 2G-H**). To confirm that there was no observable tonotopic organization in the dorsoventral axis, we compared tuning at three different depths and observed similar frequency tuning distributions (**Figure 2I**, Friedman test, p_dv_=0.4066).

### Neural selectivity, population decoding, and spatial clustering of ethologically-relevant vocalizations in the DCIC

We next sought to characterize the response properties of DCIC neurons during playback of natural vocalizations from conspecifics. Bats signal to other bats with a frequency-modulated bout ^9,28^ to claim food when competing with conspecifics (‘social’ vocalizations), while also using echolocation to track prey and steer around obstacles (‘navigation’ vocalizations)^29,30^. These vocalizations exhibit largely overlapping spectrotemporal features with subtle differences, such as shallower FM sweep rate and greater low frequency energy of social calls compared to navigation calls ^30^ (**Figure 3A**). The vast majority of recorded neurons responded to conspecific vocalizations (86±18.4%, ANOVA for responsivity, p<0.05, **Figure S4A**) whose spectral content spans from 22 to 90 kHz, highlighting a discrepancy between responsivity to vocalizations versus pure tones: in this population, neurons’ BF’s were generally below 32 kHz (**Figure 2C**). The population sound-evoked response was highest for conspecific vocalizations and lowest for pure tones (**Figure 3B**). These data suggest that ethologically relevant vocalizations drive DCIC neurons more than pure tones or even heterospecific complex stimuli, such as, mouse ultrasonic vocalizations (‘mouse USV’, **Figure 3A**). This result is consistent with electrophysiological recordings in a number of species showing that conspecific vocalizations elicit larger evoked responses than more artificial stimuli in the IC^31^. We observed a striking selectivity within individual neurons for categories of bat vocalizations (social: 30.2±4.5%; navigation: 19.7±3.7%) but little to no selectivity for mouse USVs (**Figure 3D**, p=0.13, KW test vs. pure tones, detailed selectivity in **Figure S4B**), highlighting the importance of ethological relevance in auditory response selectivity. Given the lack of selectivity to mouse USVs, we focused on the bat vocalizations for further analysis. We calculated the neural selectivity to social versus navigation calls by computing the difference in evoked responses between both categories (**Figure 3C-E, and S4C-D**). While there were neurons selective to both bat vocalization categories (social or navigation), the DCIC network exhibited a strong preference for social calls (**Figure 3E and S4D**, overall, n=1,383 social-selective neurons, n=502 navigation-selective neurons; per bat, 24±5.4% social-selective and 8.4±1.9% navigation-selective, KW test, p=5.3e-05)^30^. These data demonstrate that individual DCIC neurons are selective to conspecific vocalizations with a strong preference for encoding social vocalizations.

**Figure 3.**
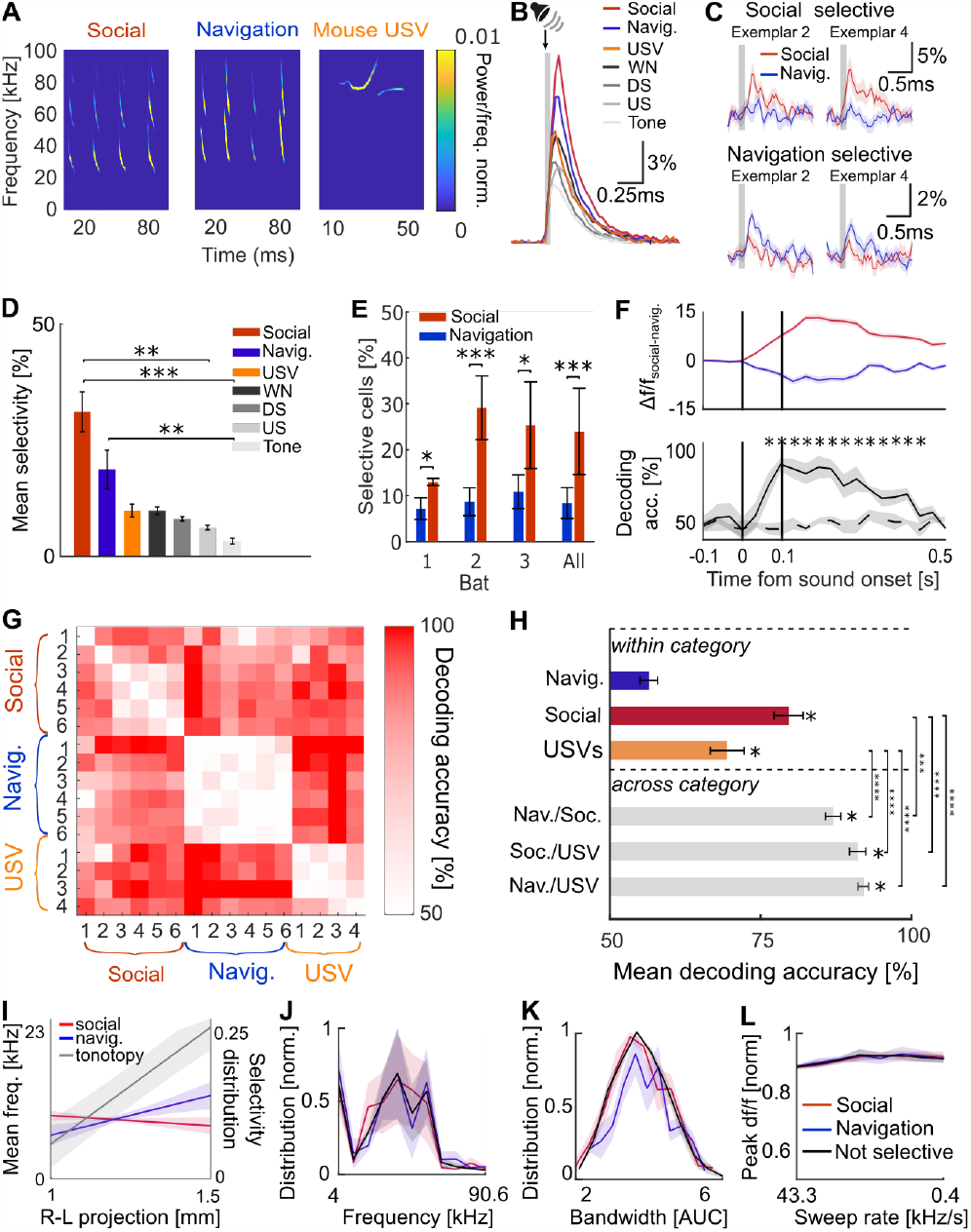
Categorical encoding of bat vocalizations in the DCIC. **A**. Example spectrograms for the vocalization categories presented. Left: social call sequence, center: temporally matched navigation call sequence, right: mouse ultrasonic vocalization (USV). **B**. Population sound-evoked response as Δf/f for all stimulus types presented (n_bat_=2, nc_ell_=2232). Conspecific bat calls evoked the highest evoked-response followed by mouse USVs and white noise, with tones evoking the smallest. All stimuli evoked significantly different responses (Friedman test on peak amplitude, α<0.05) apart from USVs and navigation calls. **C**. Example cells’ sound-evoked responses (Δf/f, median, shading: SEM) for two exemplars of social calls (red) and corresponding navigation calls (blue). Top: example social selective cell. Bottom: example navigation selective cell. **D**. Mean population selectivity for stimulus categories. The imaged population is more selective to the bat calls than other stimulus types (KW test, p_social/tone_=0.0001, p_navig.l/tone_=0.0049, p_social/us_=0.0146). **E**. Average percentage of selective cells per bat and across bats (error bar: STD). Neuronal population shows a bias towards social selectivity (KW test, p_bat1_ = 0.0495, p_bat2_ = 0.0008, p_bat3_ = 0.0495, p_all_ = 5.2683e-05). **F**. Time course of population selectivity. Call sequence onset and offset is indicated by the black lines. *Top*: difference between mean social-evoked and navigation Δf/f for top 5% social selective cells (in red, average across bats, shading: SEM) and top 5% navigation selective cells (in blue). Note that the difference in activity between social and navigation preferring cells increases rapidly after stimulus onset. *Bottom*: Time course of decoding accuracy (social vs navigation, mean across bats = plain line, shading = SEM), star indicates significance compared to chance level decoding (dashed line, permutation test, α<0.05). Bat call category can be decoded from neuronal population 60ms after sound onset before the end of the call sequence. **G**. Pairwise decoding accuracy averaged across bats for each call (conspecific or heterospecific) exemplar pairs. Decoding accuracy within categories is lower than across category (white = chance, red = 100% accuracy). **H**. Mean decoding accuracy within and across categories (error bar: SEM). Higher than chance decoding accuracy is indicated by a single star above each bar. Decoding accuracy is significantly lower vs. across categories (ANOVA, p_S/N/S_=0.0001, p_S/S/U_<e^-4^, p_U/N/S_<e^-4^, and p_U/S/U_<e^-4^). Note that decoding accuracy is not significantly different between groups within (ANOVA, p_S/U_= 0.0808) and groups across (ANOVA, p_N/S/S/U_ = 0.4619, p_N/S/N/U_ = 0.2436, p_S/U/N/U_ = 0.9999). **I**. Linear fit of cell ratio along the tonotopic gradient (gray) for both social selective (red) and navigation selective (blue). Averaged cell ratio for both social and navigation selective cells are significantly different than the tonotopic gradient (ANOVA, p <e^-4^) but not from one another (ANOVA, p = 0.9979). **J**. Mean best frequency normalized distribution for social, navigation and non-selective cells (shading: SEM, Friedman, p = 0.9688). **K**. Mean bandwidth normalized distribution for social, navigation and non-selective cells (shading: SEM, ANOVA, p = 0.8182). **L**. Mean downsweep rate tuning as peak Δf/f normalized for social, navigation and non-selective cells (shading: SEM, ANOVA, p = 0.9425).

Bat vocalizations tend to be grouped in sequences, for example the social calls presented here contain 3 to 4 successive calls. Therefore, categorical information could be carried not only by the spectral content but also by the temporal pattern of call sequences. To determine if the selectivity observed is due to the processing of the spectral information of the first call, the processing of the entire sequence or a mixture of both, we analyzed the time course of the neural selectivity. The activity of selective neurons diverged rapidly after the sound onset (**Figure 3F**, top panel) and before the end of the call sequence. A linear decoder reliably discriminated between conspecific categories on the second imaging frame after sound onset (∼32 ms, 31.25 Hz acquisition rate, **Figure 3F**, bottom, **Figure S5B** for comparison with SI), arguing that the selectivity and population-level representations arise within one to two calls.

The high proportion of social-selective neurons and rapid population discriminability suggests that the DCIC network may encode social vocalizations with high fidelity. To test this, we shifted from single-neuron analyses to population decoding. We trained a linear decoder for all pair-wise stimuli presented to the bats and found that decoding within category performed best for social vocalizations (**Figure 3G-H, Figure S5C** for comparison with SI), suggesting that the higher proportion of social-selective neurons allows the network to encode these vocalizations with higher fidelity (even if the spectral distance within social calls is lower than within mouse USVs, **Figure S5A**). We then sought to test the extent to which the DCIC network encoded call category by looking at decoding strength within versus across categories. Interestingly, decoding accuracy was significantly higher across than within categories for all category types (**Figure 3G-H**, ANOVA, p<10e^-5^ and p = 0.0001). Despite the lack of single neuron selectivity to mouse USV’s, the DCIC network reliably decoded individual calls within the mouse USV category (70±2.5%, **Figure 3G**), but to a lesser extent than across categories of bat vocalizations (91±1.4%, **Figure 3G**) which was similar to conspecific categorical decoding (**Figure 3F**, 87±1.4%, 91.1±1.4%, and 91.9±0.9% respectively, ANOVA, p_con/social/USVs_ = 0.4619, p_con/echo/USVs_ = 0.2436). The similarity in cross-category decoding between conspecific vocalizations (social vs. navigation) and conspecific vs. heterospecific (social vs. mouse UV, navigation vs. mouse USV), is at odds with the sounds’ spectral distances (**Figure S5A**) and indicates that the DCIC network contains a higher-fidelity representation of conspecific calls, likely relying on species-specific spectrotemporal filtering properties.

We next sought to understand whether neurons’ selectivity to distinct conspecific vocalizations could be explained by differences in lower-level auditory feature responses. For example, social calls have shallower frequency modulated component at the lower frequency tail of the sweep compared to navigation calls, and social-selective neurons may more strongly encode low frequency stimuli, clustering along the tonotopic axis to reflect their tuning preference. Surprisingly, this was not the case. The ratio of social-selective and navigation-selective neurons was uniformly distributed along the recorded tonotopic axis (**Figure 3I**). Importantly, other low-level features, including frequency tuning, bandwidth, duration and complex sound responsivity were identical for social and navigation selective neurons (**Figure 3J-L**, tuning: Friedman test, p = 0.9688; bandwidth, ANOVA, p = 0.8182; down sweep rate tuning, ANOVA, p = 0.9425; **Figure S6A-C**). Thus, low-level features could not explain categorical selectivity, indicating that neural selectivity for vocalizations in the DCIC may reflect a higher-order categorical representation.

Finally, we sought to evaluate whether neural encoding of vocalizations in the IC are spatially organized by exploiting the high spatial resolution of two-photon imaging. For each category-selective neuron, we computed the percentage of neighboring neurons that were also selective to the same category in 30 *μ*m steps in a site-specific manner. Surprisingly, this revealed clusters of category-selective neurons to either social or navigation calls for all bats in both hemispheres (**Figure 4**). The percentage of social selective neighboring cells is significantly higher than what would be expected by chance from a ‘spatial shuffle’ for distances of 30 *μ*m and 60 *μ*m (**Figure 4B-C**, p_30_ = 0.01, p_60_ =0.01, see inset, ANOVA, p_30-90_ = 1.2692e-07). The same relationship was observed in navigation selective cells (**Figure 4D-F**, p_30_=0.01, see inset, ANOVA, p_30-90_ = 1.2773e-04). To further describe the spatial organization of these selectivity clusters, we identified cluster centers and used a composite map to compare their location with the anatomically defined tonotopic gradient (**Figure 4D, H**). Clusters centers are distributed along the tonotopic axis and follow the cell density distribution (**Figure S7A-B** for example site and **Figure S7C** for cluster distribution). Neurons exhibiting call selectivity were, thus, spatially clustered into functional microdomains largely independent of the larger-scale tonotopic gradient.

**Figure 4.**
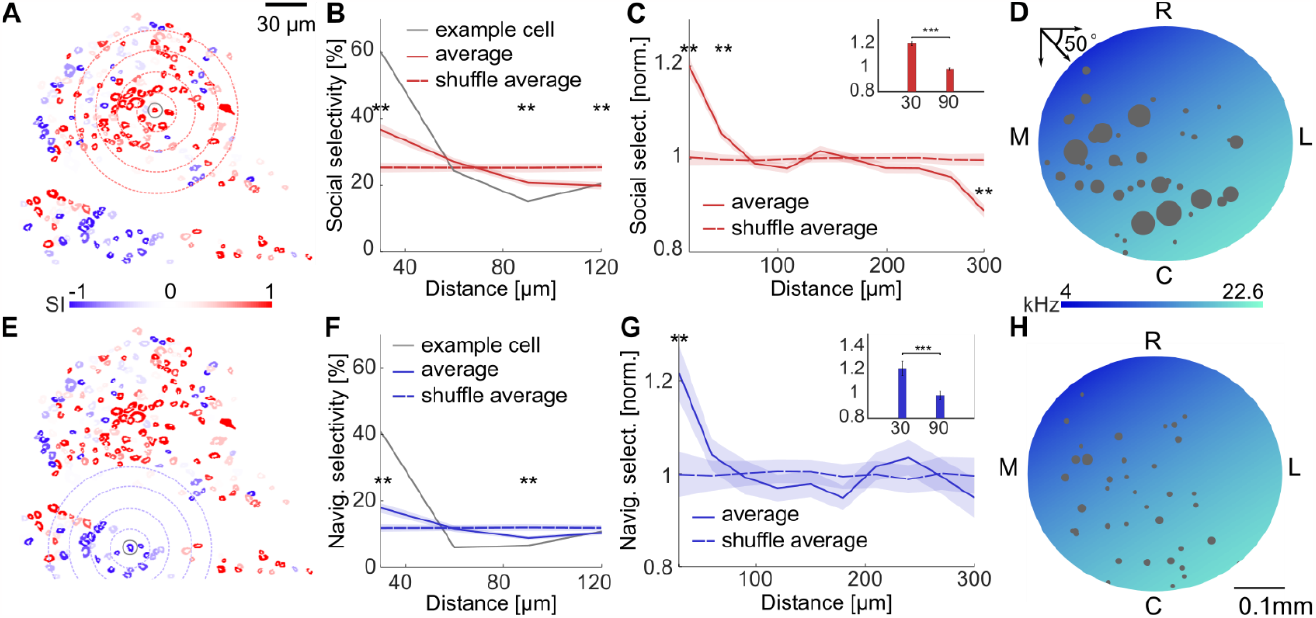
Call selectivity is spatially clustered, independent of tonotopic gradient. **A**. Example site, ROIs color-code corresponds to the social (red) vs navigation (blue) selectivity index. The social selective example cell presented in B is highlighted with a gray circle. Red dotted circles indicate areas used to compute the percentage of social selective neighboring cells (in 30 *μ*m steps). **B**. Percentage social selective cells as a function of distance from social selective cells (shading: SEM) for example site in A. Example cell highlighted in A (gray line), site average (red plain line) is significantly higher than the spatial shuffle (red dotted line) at 30 *μ*m (p = 0.01) and decreases below the shuffle line above 90 *μ*m (p_90_ = 0.01, p_120_ =0.01). **C**. Percentage of social selective cells as a function of distance from social selective cells normalized across all sites and bats. Average is higher (red plain line) than spatial shuffle (red dotted line) at distances below 60 *μ*m (p_30_ = 0.01, p_60_ =0.01) (shading: SEM) and lower at 300 *μ*m (p_300_ = 0.01). Additionally, percentage (normalized) decreases significantly between 30 and 90 *μ*m (ANOVA, p = 1.2692e-07), as displayed in the inset. **D**. Social cluster centers (gray dots, dot size is proportional to number of cells in the cluster) are distributed across composite tonotopic gradient (BF color-coded). **E**. Example site in A. The gray circle highlights navigation selective cell presented in F. Here, concentric blue dotted circles indicate the distance in 30 *μ*m steps to the example cell. **F**. Percentage of navigation selective cells as a function of distance from navigation selective cells (shading: SEM) for example site in D. Example cell highlighted in D (gray line), site average (blue plain line) is significantly higher than the spatial shuffle (blue dotted line) at 30 *μ*m (p = 0.01) and decreases below the shuffle line above 90 *μ*m (p_90_ = 0.01). **G**. Percentage of neighboring navigation selective cells as a function of distance from navigation selective cells normalized across all sites and bats. Average is higher (blue plain line) than spatial shuffle (blue dotted line) below 30 *μ*m (p_30_ = 0.01) (shading: SEM). Additionally, percentage (normalized) decreases significantly between 30 and 90 *μ*m (ANOVA, p = 1.2773e-04), as displayed in the inset. **H**. Navigation cluster centers (gray dots, dot size is proportional to number of cells) are distributed across composite tonotopic gradient (BF color-coded).

## Discussion

Current models of categorical perception rely on the idea that the periphery and midbrain serve primarily a feedforward and filter-bank role^10,32,33^. However, our data demonstrate that categorical representations of call meaning at the single-neuron and population level emerge earlier in the auditory hierarchy than posited by traditional models and display ‘hotspots’ of spatially clustered, categorical encoding. Future work will determine the extent to which this property is locally computed by the IC, at earlier stations of the brainstem, or alternatively, relies on top-down input from the neocortex. This ethologically relevant neural code for categorical information is independent of the superficial tonotopy in the DCIC. Our data support a revised view of categorical perception in which ethologically relevant sensory streams are instantiated at the single-neuron and population-level in early auditory centers to provide specified and spatially segregated acoustic channels of categorical information to recipient regions. This functional microarchitecture likely serves to increase the speed of transmission (by virtue of being early in the sensory pathway) and reduce the wiring costs (through spatially organized channels) for echolocating bats, and these computational principles likely extend to other species that extract sound meaning from rich vocal repertoires.

## Acknowledgements

We thank Dr. A. Salles for providing the conspecifics bat calls, Dr. B. Englitz for providing the mouse USVs, and C. Diebold for animal care and surgical support, Dr. Y. Boubenec for help with the spectral distance analysis. We thank Drs. C. Drieu, S. Moore, and N. Kothari for their thoughtful feedback on our manuscript. This work was supported by an NIH Brain Initiative R34 grant: R34NS118462 (KVK, CM, MW).

## Author contributions

Conceptualization: J.L., M.W., C.M., K.K.; experiments: J.L., M.W.; data analysis: J.L., K.K., writing -original draft: J.L., K.K.; writing – review & editing: J.L., M.W., C.M., K.K.

## Declaration of interests

The authors declare no competing interests.

## Materials and Methods

## Key resource table

Key resource table.
| REAGENT or RESOURCE | SOURCE | IDENTIFIER |
| --- | --- | --- |
| Bacterial and virus strains |  |  |
| pENN.AAV.CamKII, GCaMP6f.WPRE.SV40 | Addgene | 100834-AAV5 |
| Experimental models: Organisms/strains |  |  |
| *E. fuscus* | Wild caught | N/A |
| Software and algorithms |  |  |
| Matlab | The MathWorks | https://www.mathworks.com |
| ImageJ | Schneider et al.<sup>34</sup> | https://imagej.nih.gov/ij/ |

### Animals

Data was collected from three adult big brown bats (*Eptesicus fuscus*, one male and two females, ages unknown, wild caught) collected from exclusion sites in the State of Maryland. The bats were collected under the permit issued by the Maryland Department of Natural Resources (no. 55440). All procedures were approved by the Institutional Animal Care and Use Committees at Johns Hopkins University (protocol no. BA17A107), where this research was conducted.

### Anesthesia and pain management

All surgical procedures were performed under Isoflurane anesthesia (1.5%-3%) with 0.8 oxygen flow in accordance with the protocol no. BA17A107. 30 minutes prior to anesthesia induction, animals received half a dose of meloxicam (Metacam) orally for its analgesic effect. The second half dose was delivered once the animal awoke, as well as sulfamide, for antibiotic coverage. Post-surgery recovery lasted 4 to 5 days with a daily dose of meloxicam and sulfamide.

### GCaMP6f viral transfection

We injected AAV5-CamKII-GCaMP6f (Addgene, 100834-AAV5) bilaterally in the IC. We first resected the muscle located on top of the skull (*temporalis medius*) working from an incision placed above the midline to uncover the IC skull surface bilaterally. To avoid disrupting the recording surface, we entered the IC at an angle of 35° to 50° and traveled ∼1mm to reach an injection site at the center of the DCIC at a depth of -600*μ*m (see Figure 1A for visualization). We used a Hamilton syringe (10*μ*L, 700 series, 34 gauge) preloaded with the virus (1*μ*L per hemisphere) and injected using a syringe pump (Harvard Apparatus) at 75nL/min and a force of 30%. We kept the needle in place for an additional 5 minutes after injection before slowly removing from the brain. Animals were allowed to recover for at least 1 month before the next procedure (to allow for viral expression).

### Headpost implantation

We resected the muscle on the rostral part of the skull. A custom built flat two-pronged headpost (**Figure 1A**) was cemented with dental cement (Metabond) to the cranium at a distance of at least 4mm from the IC center, identified visually. The bat was then put on rest for at least one day before the thinned-skull procedure to allow for the cement to cure.

### Thinned-skull approach

We adapted thinned-skulled approached developed in the mouse ^35,36^. Using a surgical drill, we carefully thinned the skull above both hemispheres of the IC, without piercing the skull layer. The skull was then dried, and a thin layer of transparent fast-drying adhesive was applied above the thinned area. Bats were allowed to recover for a day prior to data collection.

### Two-photon calcium imaging acquisition

Two-photon calcium imaging was performed using a two-photon resonant microscope (Neurolabware) equipped with a 16X objective (Nikon). Two-photon fluorescence was excited at 980nm using an Insight X3 laser (SpectraPhysics). Data contained in this manuscript were collected at 31.25Hz over one plane at either 4X (294 μm x 192 μm, cropped) or 2X (568 μm x 374 μm, cropped). Additionally, a volumetric stack (z-stack) was collected at the end of each session in 2 or 5 *μ*m steps from the skull surface up to 200 *μ*m below the surface.

### Two-photon calcium imaging preprocessing

We used suite2p ^37^ to register and align each session. We drew regions of interest (ROIs) on the session mean image using ImageJ, as automatic ROI identification was perturbed by large differences in brightness related to changes in skull thickness over the FOV. Fluorescent time traces of individual ROIs were extracted using a custom MATLAB-based toolbox. Δf/f was computed per ROI and sound presentation by dividing the raw fluorescence by the median fluorescence for a baseline period defined as the 10 frames (312.5ms) that immediately preceded sound onset.

### Site alignment

Each imaging site position on the surface of the IC was determined by manually matching the site’s vasculature with the corresponding widefield vasculature image (larger). All widefield images collected for one hemisphere were then tiled using the vasculature as a landmark, creating a composite map per hemisphere (example of imaging sites overlay on composite widefield image in **Figure 1B**). The relative position of each imaging sites to the most anterior and medial widefield image was then computed allowing us to generate ML and RC coordinates within one hemisphere. Those coordinates were then placed in the bat’s coordinate system by manually matching the vasculature from the obtained composite maps to images covering the full extent of both ICs, obtained during the skull thinning.

### ROIs coordinates

Mediolateral and caudorostral coordinates for each cell were obtained by aligning recording sites within hemisphere and animal (n_bat_=3, n_hemi_ = 4, n_sites_=20) to an anatomical landmark obtained from surgery images covering both hemispheres: the rostro-medial boundary between SC and IC marked by the confluence of the superior sagittal and lateral sinuses. ROI depth was obtained by estimating skull-thickness above each ROI and subtracting that value from the overall site depth from brain surface identified using the z-stack. Skull thickness was computed for each FOV as points above a 95% fluorescent boundary based on the FOV raw fluorescence distribution (skull brightness is very high, **Figure S2** illustrates this method for an example site).

### Auditory stimulation

All auditory stimuli were presented contralaterally to the imaging site using one free field electrostatic speaker (ES1, TDT) placed at 45° angle, 5cm away and at a 0° azimuth from the bat’s head. All auditory stimuli were presented to the awake passively listening bat using a high-frequency auditory signal processor (RZ6, TDT) at a 200 kHz sampling rate. Sounds were generated in TDT (pure tones, white noise, upsweep, downsweep) or were recorded as WAV files and then played back via the TDT. All stimuli were calibrated at 70dB using a GRAS microphone (GRAS 46DP), placed at the bat’s ear putative position.

### Auditory stimuli

A series of stimuli were presented: pure tones ranging from 4 to 90.51kHz, 1/2 octave spaced (10 tones, pseudo random presentation, durations: 2, 4, 6, 10, 20ms), white noise, upsweep and downsweep (4khz to 90.51kHz, durations: 2, 4, 6, 10, 20, 50, 100, 200ms), bat and mouse vocalizations (5 or 6 exemplars: food claiming call sequences termed FMBs ^9,28,30^, 5 or 6 exemplars: temporally matched echolocation call sequence^30^, 2 or 4 mouse ultrasonic vocalizations^38^), and bat hunting navigation call sequences ^39^. Each sound was typically repeated 10 times (1 out of 23 analyzed sessions had 5 repetitions) and the inter-sound interval was kept constant at 1.2s for all stimuli except for the longer hunting navigation sequences where it was 2.4s.

### Histology

At the end of the experiment, the bats were perfused transcardially with 15mL of 1% phosphate buffered saline (PBS) followed by 20mL of 4% paraformaldehyde in PBS 1%. The fixed part of the brain containing the IC was sectioned in 50 *μ*m slices coronally. To ascertain excitatory expression of GCaMP6f viral expression, immunochemistry was performed following a double staining procedure for GFP and GAD67^40^.

### Data analysis

All data analysis and statistical testing were performed using custom functions written in MATLAB 2018b or 2021b (The MathWorks). Normality of data was assessed prior to statistical testing using a one-sample Kolmogorov-Smirnov test. ANOVAs were performed when the distribution was normal, while non-parametric tests were applied (Kruskalwallis or Friedman) when the data did not follow the normal distribution. The tests used to assess significance are specified in the main text and in the figure legends.

### Best Frequency identification

Best frequency (BF) was defined as the sound frequency evoking the highest response measured as the median of the peak amplitude across 50 trials (10 frames from stimulus onset, 312.5ms) at the sound level presented (70dB). BF analysis was restricted to neurons that exhibited significant tone-evoked activity (paired ttest, α<0.05).

### Tonotopic gradient characterization

We first established relationship between three anatomically defined axis rostrocaudal, mediolateral and dorsoventral (RC, ML and DV, respectively) and best frequency tuning (limited to the 4-22.7kHz range that comprises most of the tuning, **Figure 2C**) per hemisphere and animal, using a linear regression. We then computed the angle that explained tuning best by reducing the mean square error per hemisphere and bat (**Table S1**). The angle used for the rest of the analysis is the bat averaged angle 48°. The composite theoretical tonotopic gradient (**Figure 4D** and **Figure 4H**) was obtained using the averaged angle and averaged linear regression coefficients across bats.

### Sound selectivity

An individual cell is referred to as selective if its response shows invariance to members of one category while showing different responses to a second category^30,41^. We computed the selectivity index for each cell as:

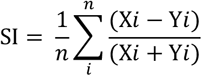

with X*i* as the median activity evoked within the response window (10 frames after sound onset, 312.5ms) by exemplar *i* of a stimulus type and Y*i* the corresponding activity evoked by an exemplar *i* of a second stimulus. Because multiple exemplars were presented, SI corresponds to the average of the selectivity index computed for pairs of exemplars. SI ranged from 1 to -1 with positive SI indicating selectivity to the first category, while negative SI indicating selectivity to the second category. Permutation tests were performed to assess cell selectivity significance (two-sided, α<0.05).

### Category decoding

We evaluated the accuracy with which the population activity could be classified as a natural stimulus exemplar by training and testing a linear discriminant classifier ^42^ in a pairwise fashion, using a leave-one-out cross-validation. Two training population vectors (*c*_*s1*_, *c*_*s2*_) were obtained by averaging the activity for 320ms (10 frames) after stimulus onset for each stimulus pair and each ROIs for a subset of randomly selected repetitions. The decoding vector *w* was defined as:

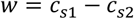

and the bias as:

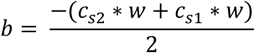

The test population vector (remaining repetition, *x*) was classified as stimulus 1 or 2, following rule:

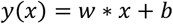

If *y(x)*>0, x is classified as stimulus 1 and as stimulus 2 if *y(x)*<0.

We repeated this procedure 100 times and averaged the accuracy computed for each stimulus pair per bat. The accuracy presented in **Figure 3G** and **S5A** is averaged across bats. Similarly, for **Figure 3F** we evaluated the accuracy with which the population could discriminate between the two bat calls categories using all exemplars of each category per time point (corresponding to the timing of a frame) and the same approach.

The weights presented in **Figure S5B-C** correspond to the average vector *w* over 100 cross-validation.

### Basic auditory feature characterization

To determine if basic functional auditory features could underlie the population selectivity to natural bat calls, we compared a number of auditory functional features for the selective and non-selective populations, including BF tuning, tuning curve bandwidth, amplitude of evoke-response to various sounds, upsweed/downsweep tuning (**Figure 3I-L** and **Figure S6**). Bandwidth is defined as the area under the curve for a normalized tuning curve (small if the tuning is sharp and large if the tuning is broad). Population tuning to downseep, upsweep rate and white noise duration was computed as the average of the tuning curves normalized by the peak amplitude.

### Spatial clustering analysis

Local percentage of selective cells was computed in 30*μ*m radial increments for each selective cell for either social or navigation selectivity per recording sites. Given the variations in site selectivity the local percentage was divided by the site overall selectivity to compute a local selectivity ratio as a function of distance per bat. Spatial shuffle distributions were computed per 30*μ*m radial increments and imaging site using the x and y coordinates (unpaired) to determine significance using a permutation test (two-sided, α<0.05). Selective clusters were identified on a composite map across sites and bats using Density-Based Spatial Clustering of Applications with Noise (*DBSCAN*, based on the pairwise Euclidean distance between ROIs). Clusters minimum number of cells was set to 3 and epsilon was set to 30*μ*m.

## Supplementary Materials

**Figure S1.**
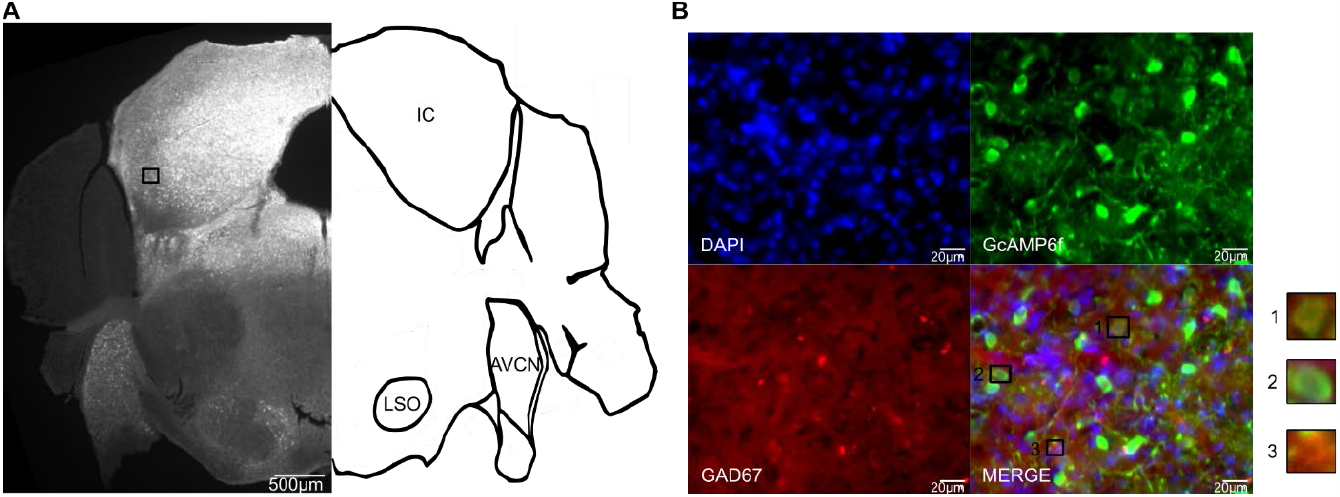
Histology and immunostaining for GCaMP6f and GAD67. **A**. GFP immunostaining shows expression of GCaMP6f throughout the IC. The black box indicates the location of the site highlighted in B. **B**. Immunostaining for IC locus highlighted in A for cell nucleus (DAPI, top left), GCaMP6f (GFP, top right), inhibitory neurons (GAD67, bottom left), and merged image (bottom right). Merged image suggests minimal overlap between GCcAMP6f and GAD67 expression. Example neurons numbered on the merged image are displayed on the right: 1) GAD67+ and GcAMP6f+ (yellow), 2) GcAMP6f+ and GAD67-(green), 3) GcAMP6f- and GAD67+ (red).

**Figure S2.**
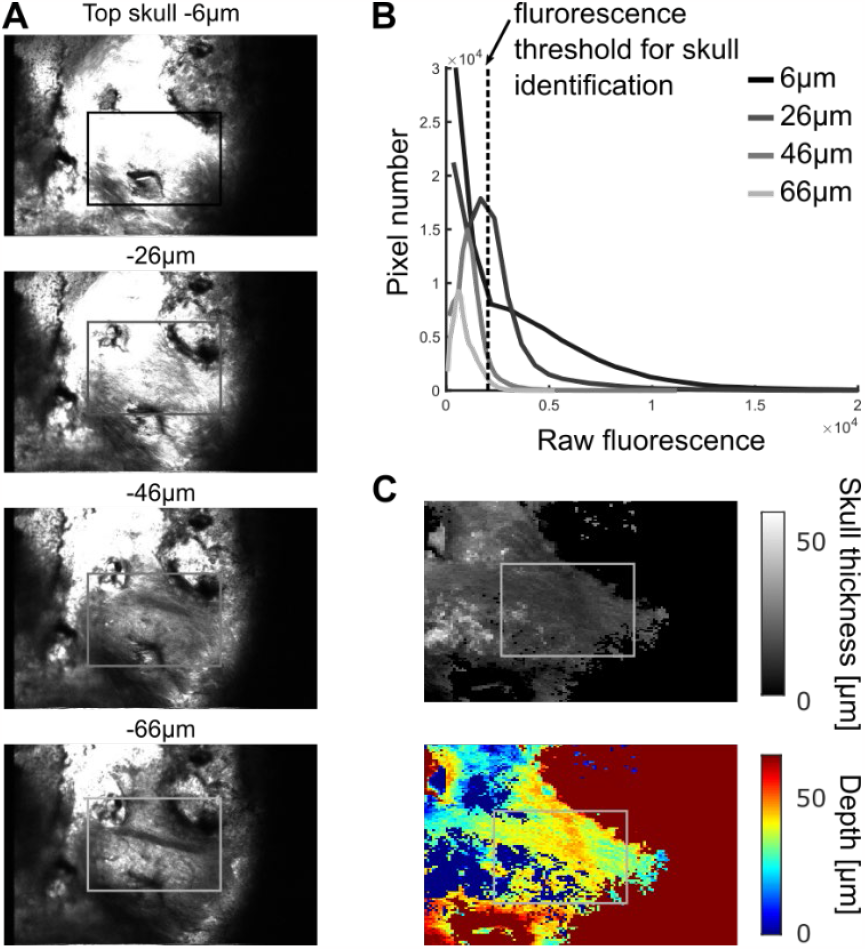
Site depth estimation corrected by skull thickness. **A**. Mean images (150 frames) from 1X volumetric stack for example 2X imaging site (highlighted by the black to gray box, colors correspond to depth in B) every 10 *μ*m until example site imaging depth (bottom image). **B**. Raw fluorescence distribution for images displayed in A. Note that shallower images possess a longer high fluorescence tail indicating a larger skull coverage. The criterion used for skull identification (95th percentile of the overall volumetric stack distribution) is displayed in the dashed line. **C**. Skull thickness and corresponding depth estimation for example volumetric stack in A and B.

**Figure S3.**
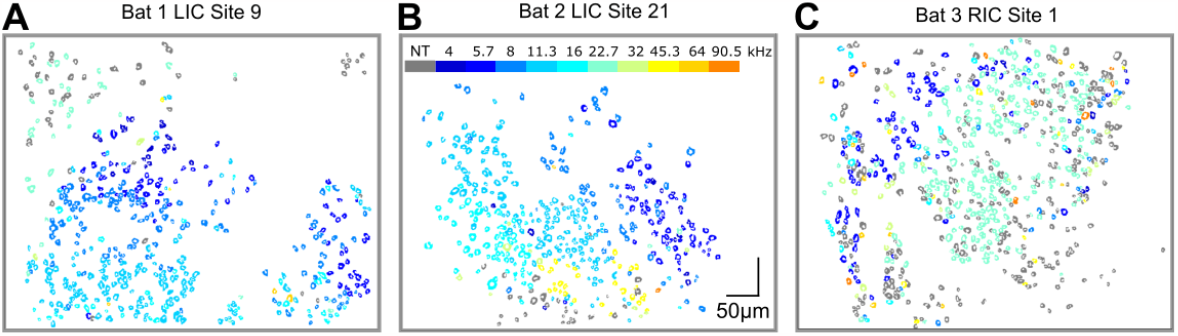
Tonotopic gradient in example sites for 3 separate bats. **A**. Example site from the left DCIC of bat 1. **B**. Example site from the left DCIC of bat 2. **C**. Example site from the right DCIC of bat 3.

**Figure S4.**
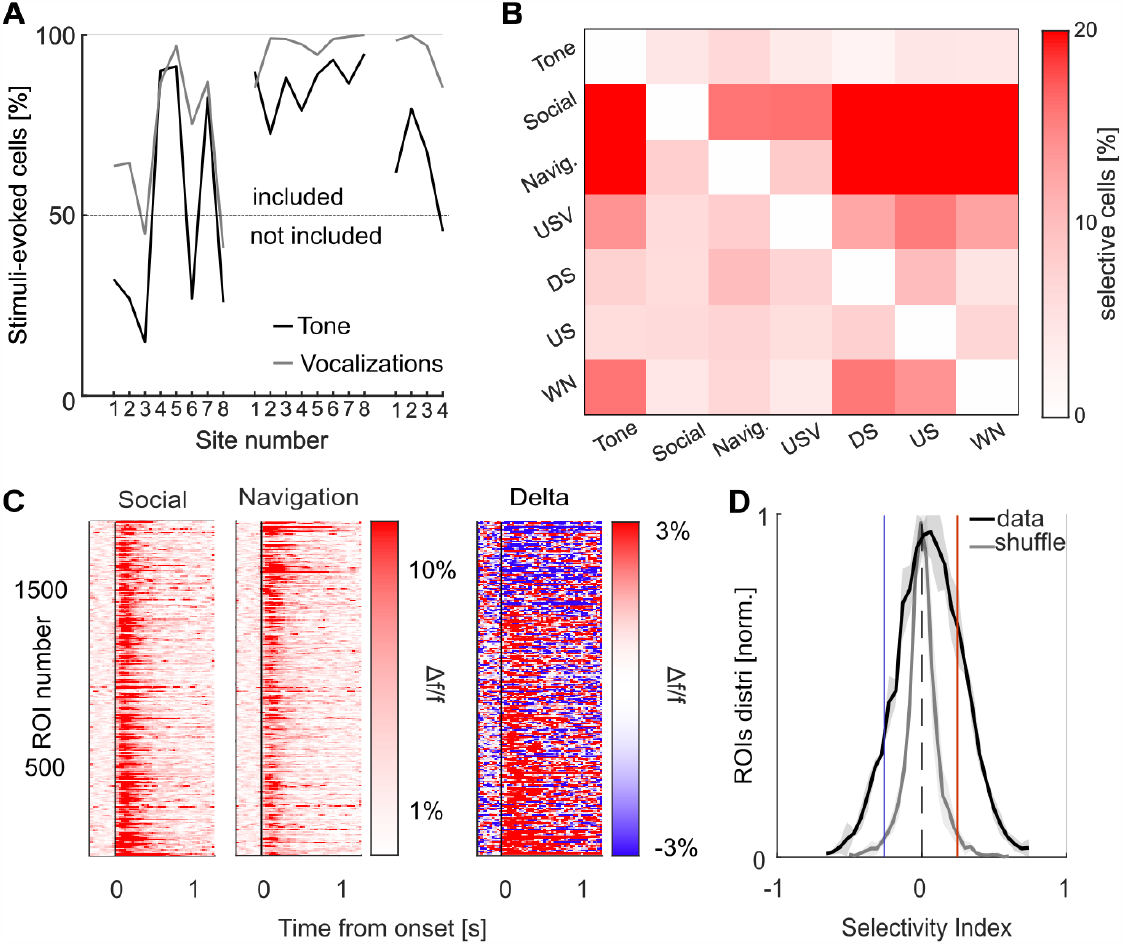
Selectivity index method. **A**. Percentage of cells showing significant stimulus-evoked responses (ANOVA, α<0.05) for tones (in black) and vocalizations (in gray) for each recorded site. Sites with fewer than 50% of cells with significant tone-evoked responses were excluded from the vocalization analysis. **B**. Mean percentage of significantly selective cells using SI for each sound category. Note that the percentage of selective cells to conspecific calls is consistently higher than other categories. **C**. Average sound-evoked Δf/f per category for significant selective cells sorted by their selectivity index (n_cells_ = 1885). Left: average sound-evoked activity for social calls. Middle: average sound-evoked activity for navigation calls. Right: difference between social and navigation activity. **D**. Average selectivity index distribution across bats (shading: SEM). Average significance boundaries are displayed in blue (2.5 percentile, SI_bound_ = -0.2959) and red (97.5 percentile, SI_bound_ = 0.3109).

**Figure S5.**
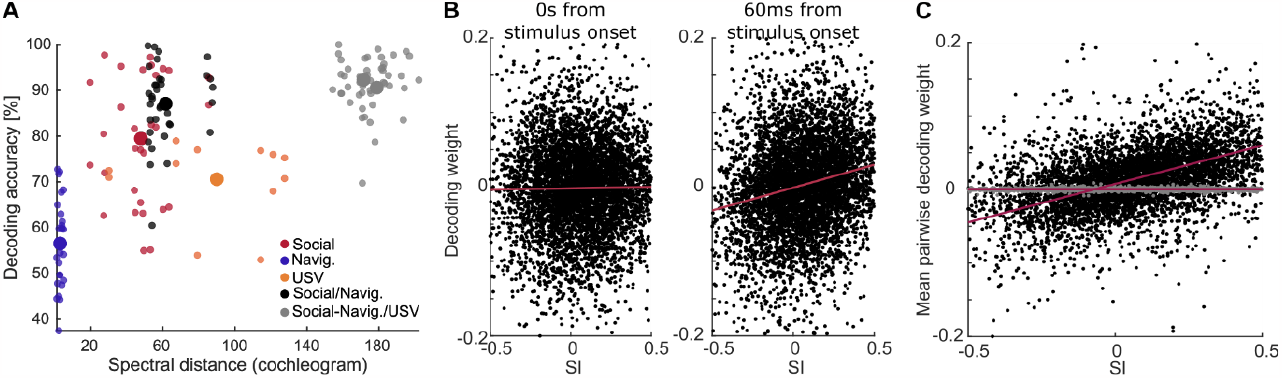
Linear decoding method. **A**. Pairwise mean decoding accuracy as a function of spectral distance for social calls (in red), navigation calls (in blue), USVs (orange), conspecific calls (social vs navigation), or cross-species calls (social vs USVs, and navigation vs USVs). Smaller dots: individual exemplars pair values, larger dots: average for one comparison type. **B**. Decoding weights for timecourse decoder (Figure 3F) as a function of selectivity index per cell (black dot). *Left*: decoding weights does not increase (linear regression in red) with SI at 0s after stimulus onset. *Right*: decoding weights increases linearly with SI at 60ms after stimulus onset. **C**. Mean decoding weights from pairwise decoding for social vs navigation (Figure 3G-H) increase with SI (linear regression in red), but not for same stimulus decoding (social only, in grey)

**Figure S6.**
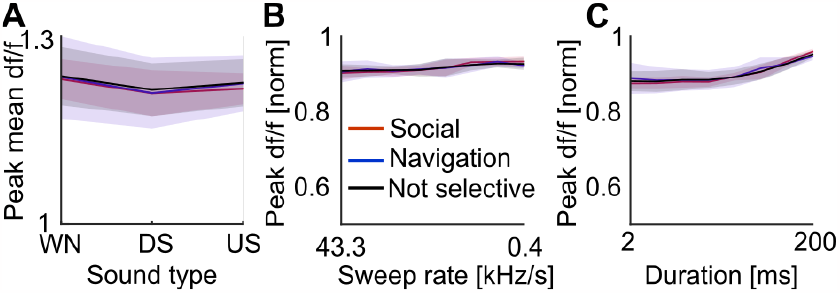
Additional auditory features for selective and non-selective populations. **A**. Average peak evoked-responses as df/f to different type of complex sounds: white noise (WN), downsweep (DS) and upsweep (US), for social, navigation and non-selective cells (shading: SEM, ANOVA, p = 0.9721). **B**. Mean upsweep rate tuning as peak df/f normalized for social, navigation and non-selective cells (shading: SEM, ANOVA, p = 0.9907). **C**. Mean white noise duration tuning as peak df/f normalized for social, navigation and non-selective cells (shading: SEM, ANOVA, p =0.9648).

**Figure S7.**
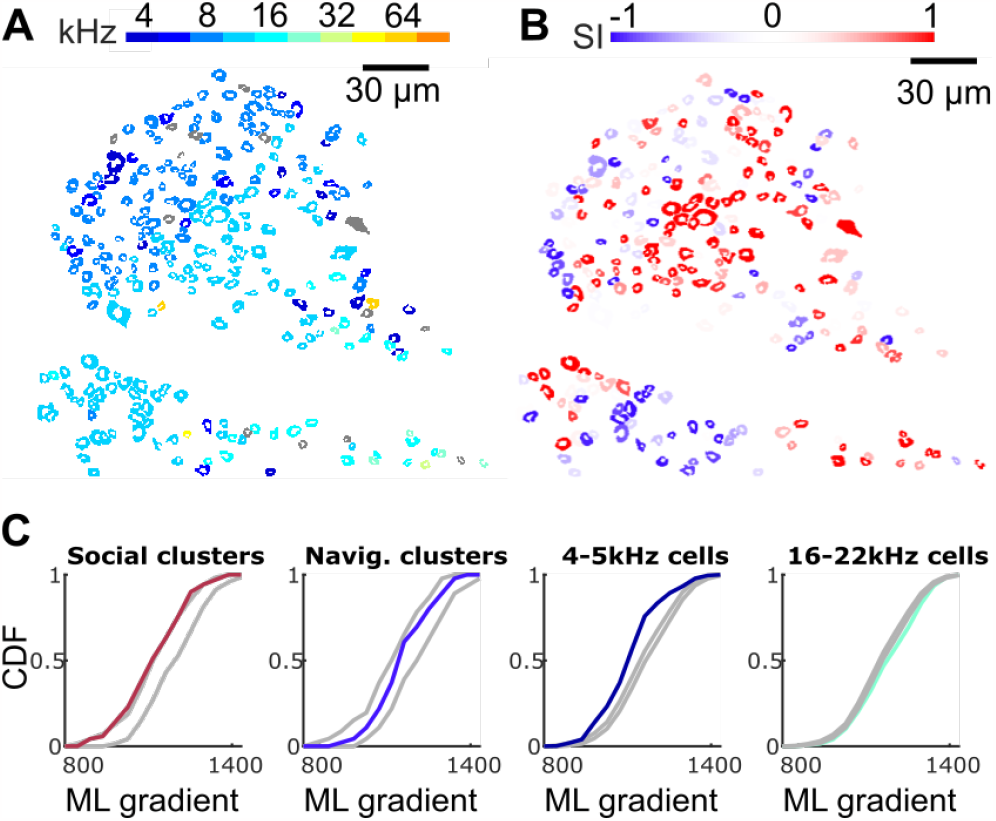
Relationship between tuning and selectivity index examples and composite view. **A**. Tuning map for example site in Figure 4 color-coded according to the best frequency of the neuron. **B**. Corresponding social/navigation selectivity map for the same example site, color-coded by the neuron’s selectivity index. Note that the selectivity ‘hotspots’ do not follow the tonotopic gradient. **C**. Cumulative distribution for cluster centers and corresponding shuffle distribution (randomly selected from all cells coordinates, in gray) along the composite RL gradient of all imaged sites. *Left*: social cluster falls just outside of the shuffle distribution (p<0.01), perhaps due to the increase in cluster size with cell density. *Center left*: navigation cluster centers fall within the 95% confidence interval therefore following cell density independently of the tonotopic gradient. *Center right*: cells tuned to low frequencies (4 and 5.7kHz, in dark blue) are located more rostromedially than the corresponding shuffled distribution (p<0.01). *Right*: cells tuned to high frequencies (16 and 22.7kHz) are located more caudolaterally than the corresponding shuffled distribution (p<0.01).

**Table S1.**
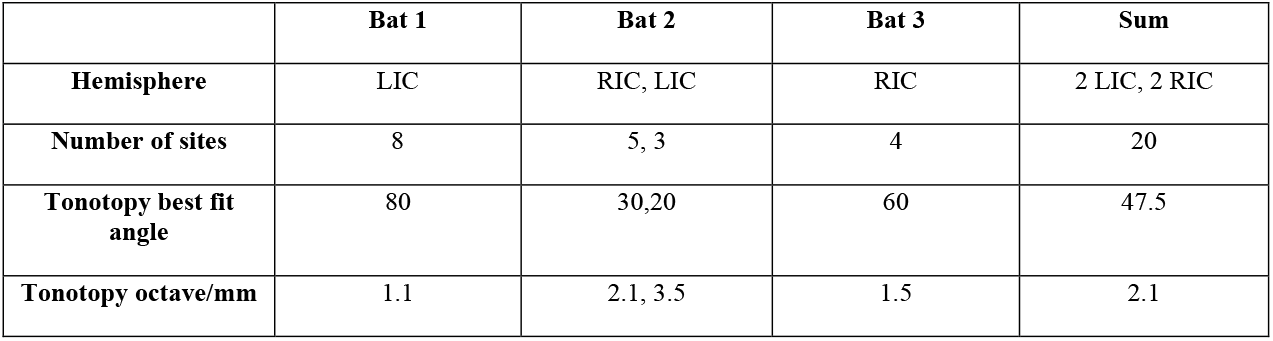
Tonotopic gradient angle estimation.

|  | **Bat 1** | **Bat 2** | **Bat 3** | **Sum** |
| --- | --- | --- | --- | --- |
| **Hemisphere** | LIC | RIC, LIC | RIC | 2 LIC, 2 RIC |
| **Number of sites** | 8 | 5, 3 | 4 | 20 |
| **Tonotopy best fit angle** | 80 | 30,20 | 60 | 47.5 |
| **Tonotopy octave/mm** | 1.1 | 2.1, 3.5 | 1.5 | 2.1 |

## Notes

### Competing Interest Statement

The authors have declared no competing interest.

